# Selective inhibition of HDAC3 targets synthetic vulnerabilities and activates immune surveillance in lymphoma

**DOI:** 10.1101/531954

**Authors:** Patrizia Mondello, Saber Tadros, Matt Teater, Lorena Fontan, Aaron Y. Chang, Neeraj Jain, Shailbala Singh, Man Chun John Ma, Haopeng Yang, Eneda Toska, Stefan Alig, Matthew Durant, Elisa de Stanchina, Anja Mottok, Loretta Nastoupil, Sattva Neelapu, Oliver Weigert, Giorgio Inghirami, Josè Baselga, Ahmet Dhogan, Anas Younes, Cassian Yee, David A. Scheinberg, Ari Melnick, Michael R. Green

## Abstract

*CREBBP* mutations are highly recurrent in B-cell lymphomas and either inactivate its histone acetyltransferase (HAT) domain or truncate the protein. Herein, we show that these two classes of mutations yield different degrees of disruption of the epigenome, with HAT mutations being more severe and associated with inferior clinical outcome. Genes perturbed by *CREBBP* mutation are direct targets of the BCL6/HDAC3 onco-repressor complex. Accordingly, we show that HDAC3 selective inhibitors fully reverse *CREBBP* mutant aberrant epigenetic programming resulting in: a) growth inhibition of lymphoma cells through induction of BCL6 target genes such as *CDKN1A* and b) restoration of immune surveillance due to induction of BCL6 repressed IFN pathway and antigen presentation genes. By reactivating these genes, exposure to HDAC3-i restored the ability of tumor infiltrating lymphocytes to kill DLBCL cells in an MHC II and MHC I dependent manner. Hence HDAC3-i represent a novel mechanism-based immune-epigenetic therapy for *CREBBP* mutant lymphomas.

**STATEMENT OF SIGNIFICANCE:** We have leveraged the molecular characterization of different types of *CREBBP* mutations to define a rational approach for targeting these mutations through selective inhibition of HDAC3. This represents an attractive therapeutic avenue for targeting synthetic vulnerabilities in *CREBBP* mutant cells in tandem with promoting anti-tumor immunity.

## INTRODUCTION

Diffuse large B-cell lymphoma (DLBCL) and follicular lymphoma (FL) are the two most frequent subtypes of non-Hodgkin lymphoma. These diseases originate from germinal center B (GCB)-cells; a stage of development that naturally allows for the proliferation and affinity maturation of antigen-experienced B-cells to produce terminally-differentiated memory B-cells or plasma cells. The germinal center (GC) reaction is regulated by B-cell-intrinsic activation and suppression of large sets of genes by master regulators such as the BCL6 transcription factor^1^, and extrinsically via the interaction of GCB-cells with follicular helper T (T_FH_)-cells and other immune cells within the GC^2^. The BCL6 transcription factor is critical for GCB-cell development and coordinately suppresses the expression of large sets of genes by recruiting SMRT and NCOR co-repressor complexes containing HDAC3^3^ and tethering a non-canonical polycomb repressor 1-like complex in cooperation with EZH2^4^. These genes are normally reactivated to drive GC exit and terminal differentiation, but the epigenetic control of these dynamically-regulated GC transcriptional programs is perturbed in DLBCL and FL via the downstream effects of somatic mutation of chromatin modifying genes^5^ (CMG).

The second most frequently mutated CMG in both DLBCL and FL is the *CREBBP* gene, which encodes a histone acetyltransferase that activates transcription via acetylation of histone H3 lysine 27 (H3K27Ac) and other residues. We have previously found that these mutations arise as early events during the genomic evolution of FL and reside in a population of tumor repopulating cells, often referred to as common progenitor cells (CPCs)^6^. We have also noted an association between *CREBBP* inactivation and reduced expression of MHC class II in human and murine lymphomas^6,7^. The expression of MHC class II is critical for the terminal differentiation of B-cells through the GC reaction^8^. The interaction with helper T-cells via MHC class II results in B-cell co-stimulation through CD40 that drives NFκB activation and subsequent IRF4-driven suppression of BCL6. However, in B-cell lymphoma, tumor antigens may also be presented in MHC class II and recognized by CD4 T-cells that drive an anti-tumor immune response^9,10^. The active suppression of MHC class II expression in B-cell lymphoma may therefore be driven by evolutionary pressure against MHC class II-binding tumor antigens, in line with that recognized in other cancers^11^. In support of this notion, the reduced expression of MHC class II has been found to be associated with poor outcome in DLBCL^12^.

Recently, MHC class II expression has been defined as an important component of interferon-gamma (IFN-γ) related signatures that are predictive of the activity of PD-1 neutralizing antibodies^13-15^. This is consistent with a prominent role for CD4 T-cells in directing anti-tumor immunity and responses to immunotherapy^16^. Despite this, current immunotherapeutic strategies largely rely on the pre-existence of an inflammatory microenvironment for therapeutic efficacy. Here, we have characterized the molecular consequences of *CREBBP* mutations and identified BCL6-regulated cell cycle, differentiation, and IFN signaling pathways as core features that are aberrantly silenced at the epigenetic and transcriptional level. We show that HDAC3 inhibition specifically restores these pathways thus suppressing growth and most critically enabling T-cells to recognize and kill lymphoma cells. Together, these highlight multiple mechanisms by which selective inhibition of HDAC3 can drive tumor-intrinsic killing as well as activate IFN-γ signaling and anti-tumor immunity which extends to both *CREBBP* wild-type and *CREBBP* mutant tumors.

## MATERIALS AND METHODS

For detailed methodology, please refer to the supplementary methods section.

### CRISPR/Cas9-modification of lymphoma cells

The RL cell-line (ATCC CRL-2261) was modified by electroporation of one of two unique gRNA sequences in the pSpCas9(BB)-2A-GFP vector (Addgene plasmid #48138, gift from Feng Zhang)^17^ with a single-stranded oligonucleotide donor template. Single GFP-positive cells were sorted 3-4 days after transfection, colonies expanded and evaluated for changes in the targeted region using Sanger Sequencing. The process was repeated for until point mutants were retrieved from each of the two gRNAs, totaling 742 single clones. All cells were maintained as sub-confluent culture in RPMI medium with 10% FBS and PenStrep and re-validated by Sanger sequencing prior to each set of experiments.

### ChIP-sequencing

Cells were washed and fixed in formaldehyde, and chromatin sheared by sonication. An antibody specific to H3K27Ac (Active Motif) was coupled to magnetic protein G beads, incubated with chromatin overnight, and immunoprecipitation performed. Input controls were reserved for comparison. Nucleosomal DNA was isolated and either used as a template for qPCR (ChIP-qPCR) or to generate NGS libraries using KAPA Hyper Prep Kits (Roche) and TruSeq adaptors (Bioo Scientific) using 6 cycles of PCR enrichment. Libraries were 6-plexed and sequenced with 2×100bp reads on a HiSeq-4000 (Illumina). The data were mapped using BWA, peaks called using MACS2, and differential analyses performed using DiffBind. For gene set enrichment analyses, the gene with the closest transcription start site to the peak was used. The statistical thresholds for significance were q<0.05 and fold-change>1.5.

### RNA-sequencing

Total RNA was isolated using AllPrep DNA/RNA kits (Qiagen) and evaluated for quality on a Tapestation 4200 instrument (Agilent). Total RNA (1µg) was used for library preparation with KAPA HyperPrep RNA kits with RiboErase (Roche) and TruSeq adapters (Bioo Scientific). Libraries were validated on a Tapestation 4200 instrument (Agilent), quantified by Qubit dsDNA kit (Life Technologies), 6-plexed, and sequenced on a HiSeq4000 instrument at the MD Anderson Sequencing and Microarray Facility using 2×100bp reads. Reads were aligned with STAR, and differential gene expression analysis performed with DEseq2. The statistical thresholds for significance were q<0.05 and fold-change>2.

### Cell proliferation assays

Cells were seeded in 96-well plates at 50,000 cell/100 μl/well with either vehicle (DMSO 0.1%) or increasing concentrations of drugs. Cell viability was assessed with the fluorescent redox dye CellTiter-Blue (Promega). The reagent was added to the culture medium at 1:5 dilution, according to manufacturer`s instructions. Procedures to determine the effects of certain conditions on cell proliferation and apoptosis were performed in 3 independent experiments. The 2-tailed Student t test and Wilcoxon Rank test were used to estimate the statistical significance of differences among results from the 3 experiments. Significance was set at P < .05. The PRISM software was used for the statistical analyses.

### Patient-derived xenograft and in vitro organoid studies

Six week old female NSG mice were implanted subcutaneously with tumor specimens from three lymphoma PDX models (NY-DR2, DANA and TONY). For the efficacy study, treatments started when tumors reached 100 mm3. Mice (12/group) were randomized and dosed via oral gavage with BRD3308 (25 mg/kg) or control vehicle (0.5% methyl cellulose, 0.2% tween 80) twice daily for 21 consecutive days. Mice were cared for in accordance with guidelines approved by the Memorial Sloan Kettering Cancer Center Institutional Animal Care and Use Committee and Research Animal Resource Center.

For in vitro organoid culture, the tumors were dissociated to single cells, stained with CFSE, washed and mixed with irradiated 40LB cells at a 10:1 ratio of Primary:40LB. The cell mixture was then used to fabricate organoids in a 96-well plate as previously described^18^, with 20 µL organoids containing 3% silicate nanoparticles and a 5% gelatin in IMDM medium solution. The organoids were cultured in IMDM medium containing 20% FBS supplemented with antibiotics and normocin (Invivogen) for 6 days, doubling the volume of medium after 3 days. The cell mixture was exposed to 4 1:3 serial dilutions of BRD3308 starting at 5 µM or vehicle control (DMSO) in triplicate for 6 days, treating a second time at 3 days. After 6 days of exposure, cell viability and proliferation were assessed by flow cytometry using DAPI staining gating on CFSE-positive cells.

## RESULTS

### *CREBBP^R1446C^* mutations function as a dominant-negative to suppress BCL6 co-regulated epigenetic and transcriptional programs

The *CREBBP* gene is predominantly targeted by point mutations that result in single amino acid substitutions within the lysine acetyltransferase (KAT) domain^6,19^, with a hotspot at arginine 1446 (R1446), which lead to a catalytically inactive protein^20,21^. However, all of the prior studies characterizing effects of *CREBBP* mutation have been performed using knock-out or knock-down of *Crebbp*, resulting in loss-of-protein (LOP)^7,19,22–24^. Furthermore, mutations of R1446 have not been documented in any lymphoma cell line. We therefore opted to investigate whether there may be unique functional consequences of KAT domain hotspot mutations of *CREBBP*. To achieve this, we utilized CRISPR/Cas9-mediated gene editing with two unique guide-RNAs (gRNA) to introduce the most common *CREBBP* mutation, R1446C, into a *CREBBP* wild-type cell line bearing the t(14;18)(q21;q32) translocation, RL (Figure 1A). This allowed us to generate clones from each gRNA that had received the constructs but remained wild-type (*CREBBP^WT^*), those that edited their genomes to introduce the point mutations into both alleles (*CREBBP^R1446C^*), and those that acquired homozygous frameshift mutations resulting in LOP (*CREBBP^KO^*) (**Figure S1**). These isogenic sets of clones differ only in their *CREBBP* mutation status, and therefore allow for detailed functional characterization in a highly controlled setting.

**Figure 1:**
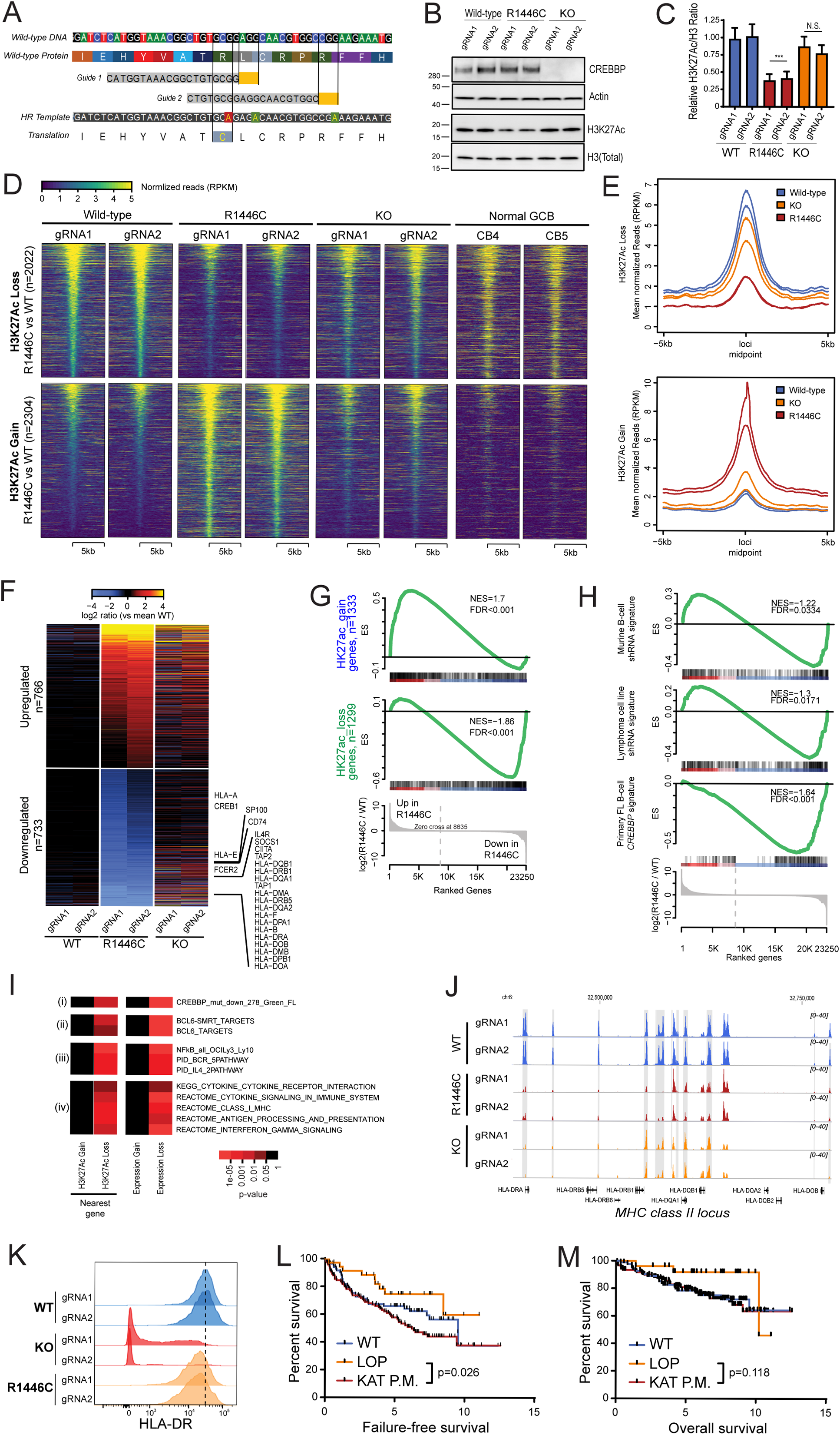
Detailed molecular characterization of CREBBP^R1446C^ and CREBBP^KO^ mutations using isogenic CRISPR/Cas9-modified lymphoma cells. **A)** A diagram shows the CRISPR/Cas9 gene editing strategy. Two guides were designed that were proximal to the R1446 codon, with PAM sites highlighted in yellow. A single stranded Homologous Recombination (HR) template was utilized that encoded silent single nucleotide changes that interfered with the PAM sites but did not change the protein coding sequence, and an additional single nucleotide change that encoded the R1446C mutation. **B)** A representative western blot shows that the CREBBPR1446C protein is expressed at similar levels to that of wild-type CREBBP, whereas CREBBPKO results in a complete loss of protein expression as expected. The level of H3K27Ac shows a more visible reduction in *CREBBP^R1446C^* cells compared to isogenic *CREBBP^WT^* cells than that observed in *CREBBP^KO^* cells. **C)** Quantification of triplicate western blot experiments shows that there is a significant reduction of H3K27Ac in *CREBBP^R1446C^* cells compared to *CREBBP^WT^* cells (T-test p-value <0.001). A reduction is also observed in *CREBBP^KO^* cells, but this was not statistically significant (T-test p-value = 0.106). **D)** Heat maps show the regions of significant H3K27Ac loss (n=2022, above) and gain (n=2304, below) in *CREBBP^R1446C^* cells compared to isogenic WT controls. The regions with reduced H3K27Ac in *CREBBP^R1446C^* cells can be seen to normally bear this mark in GCB cells. **E)** Density plots show that the degree of H3K27Ac loss (above) is most notable in *CREBBP^R1446C^* cells compared to isogenic WT cells, while *CREBBP^KO^* cells show an intermediate level of loss. Regions with H3K27Ac gain (below) in *CREBBP^R1446C^* cells are less reproducibly increased in *CREBBP^KO^* cells. **F)** A heat map of RNA-seq data shows that there are a similar number of genes with increased (n=766) and decreased (n=733) expression in *CREBBP^R1446C^* cells compared to isogenic WT controls. The *CREBBP^KO^* cells again show an intermediate level of change, with expression between that of *CREBBP^WT^* and *CREBBP^R1446C^* cells. **G)** Gene set enrichment analysis of the genes most closely associated with regions of H3K27Ac gain (above) or loss (below) shows that these epigenetic changes are significantly associated with coordinately increased or decreased expression in *CREBBP^R144C^* cells compared to isogenic WT controls, respectively. **H)** Gene set enrichment analysis shows that genes which were previously found to be down-regulated following shRNA-mediated knock-down of *CREBBP* in murine B-cells (top) or human lymphoma cell lines (middle) are also reduced in *CREBBP^R1446C^* mutant cells compared to *CREBBP^WT^* cells. However, the most significant enrichment was observed for the signature of genes that we found to be significantly reduced in primary human FL with *CREBBP* mutation compared to *CREBBP* wild-type tumors. **I)** Hypergeometric enrichment analysis identified sets of genes that were significantly over-represented in those with altered H3K27Ac or expression in *CREBBP^R1446C^* cells. This included (i) gene sets associated with CREBBP mutation in primary tumors, (ii) BCL6 target genes, (iii) BCR and IL4 signaling pathways, and (iv) gene sets involving immune responses, antigen presentation and interferon signaling were significantly enriched. **J)** ChIP-seq tracks of the MHC class II locus on chromosome 6 are shown for isogenic *CREBBP^WT^* (blue), *CREBBP^KO^* (orange), and *CREBBP^R1446C^* (red) cells, with regions of significant H3K27Ac loss shaded in grey. A significant reduction can be observed between *CREBBP^WT^* and *CREBBP^R1446C^* cells, with *CREBBP^KO^* cells harboring an intermediate level H3K27Ac over these loci. **K)** Flow cytometry for HLA-DR shows that reduced H3K27Ac over the MHC class II region is associated with changes of cell surface protein expression. A ~2-fold reduction is observed in *CREBBP^KO^* cell compared to *CREBBP^WT^*, but a dramatic ~39-fold reduction is observed in *CREBBP^R1446C^* cells. **L)** Kaplan-Meier plots show the failure free survival in 231 previously untreated FL patients according to their *CREBBP* mutation status. Nonsense/frameshift mutations that create a loss-of-protein (LOP) are associated with a significantly better failure-free survival compared to KAT domain point mutations (KAT P.M.; log-rank P=0.026). **M)** Kaplan-Meier plots show the failure free survival in 231 previously untreated FL patients according to their *CREBBP* mutation status. Patients with LOP mutations have a trend towards better overall survival, but this is not statistically significant (log-rank P=0.118).

Western blot confirmed that the CREBBP^R1446C^ protein is still expressed and that the *CREBBP^KO^* mutations resulted in a complete loss of protein expression (Figure 1B). Densitometry of the H3K27Ac mark revealed that *CREBBP^R1446C^* cells showed a significant reduction in H3K27Ac compared to isogenic *CREBBP^WT^* controls (p<0.001; Figure 1C). Although *CREBBP^KO^* cells showed levels that were lower than isogenic *CREBBP^WT^* controls, this reduction was not statistically significant (p=0.106). We therefore performed chromatin immunoprecipitation (ChIP)-sequencing for H3K27Ac to define the physical location of these changes and identify potentially deregulated genes. This revealed 2022 regions with significantly reduced acetylation, and 2304 regions with significantly increased acetylation in *CREBBP^R1446C^* cells compared to isogenic *CREBBP^WT^* controls (Fold change >1.5, Q-value<0.01; Figure 1D, **Table S1**). Regions with loss of H3K27Ac were observed to normally bear this mark in human GCB cells^24^, suggesting that *CREBBP^R1446C^* mutations lead to loss of a normal GCB epigenetic program. Notably, *CREBBP^KO^* resulted in a reduction of H3K27Ac in these regions also, but at a lower magnitude than that observed with *CREBBP^R1446C^* mutations (Figure 1D-E). This was not as consistently observed for regions with increased H3K27Ac, which showed fewer changes in *CREBBP^KO^* cells.

Using RNA sequencing, we observed broad changes in transcription, with 766 genes showing significantly increased expression and 733 genes showing significantly decreased expression in *CREBBP^R1446C^* cells compared to *CREBBP^WT^* isogenic controls (Fold change >1.5, Q-value<0.01; Figure 1F; **Table S2**). The genes that were proximal to regions of H3K27Ac loss showed a coordinate reduction in transcript abundance and *vice versa* (Figure 1G), suggesting that these broad changes in transcription are directly linked to altered promoter/enhancer activity as a result of differential H3K27Ac (52). Importantly, we observed significant enrichment of the transcriptional signature that we have previously characterized using shRNA knock-down of *CREBBP* in murine B-cells or human DLBCL cell lines^7^, with genes that are down-regulated following shRNA knock-down also being significantly reduced in *CREBBP* mutants (FDR<0.05, Figure 1H). However, a more significant enrichment was observed for the signature of genes we defined as being lost in association with *CREBBP* mutation in primary human FL B-cells^6^ (FDR<0.001, Figure 1H). This shows that these CRISPR/Cas9-edited clones closely resemble the molecular phenotype of *CREBBP* mutations in primary B-cell lymphoma, and therefore provides a tractable model for studying the functional consequences of these mutations.

To provide insight into the pathways that are deregulated by *CREBBP* mutation, we used hypergeometric analysis to identify sets of genes that were significantly deregulated by *CREBBP^R1446C^* mutations at the epigenetic and transcriptional level (Figure 1I, **Table S3-4**). This revealed a significant enrichment of BCL6-SMRT and BCL6 targets, including those with a role in BCR, NFκB, in addition to interferon signaling and antigen presentation (Figure 1I). In line with the enrichment of antigen presentation pathways and the observed genome-wide differences in H3K27Ac in *CREBBP^KO^* cells compared to *CREBBP^R1446C^* cells, we also noted a more modest reduction of H3K27Ac over the MHC class II gene locus on chromosome 6 in *CREBBP^KO^* cells (Figure 1J). This was associated with a ~2-fold reduction of MHC class II on the cell surface of *CREBBP^KO^* cells compared to isogenic *CREBBP^WT^* controls, but a ~37-fold reduction in *CREBBP^R1446C^* clones (Figure 1K). These data further support a stronger epigenetic and transcriptional suppression of IFN-responsive genes, including those involved in antigen presentation, in *CREBBP^R1446C^* mutations compared to *CREBBP^KO^*.

Mutations of *CREBBP* have been previously associated with adverse outcome in FL and are incorporated into the M7-FLIPI prognostic index^25^. However, in these analyses, all *CREBBP* mutations were considered collectively without discriminating between KAT domain point mutations or nonsense/frameshift LOP mutations. We therefore re-evaluated these data in light of our observed functional differences between these mutations. This showed that there was a significant difference in failure-free survival (Figure 1L) between these two classes of *CREBBP* mutations. This was not significant for overall survival (Figure 1M). Specifically, patients bearing KAT domain point mutations in *CREBBP* (22% of which were R1446 mutations) had a significantly reduced failure-free survival compared to patients with LOP mutations in *CREBBP* (Log-Rank P=0.026).

These results provide the first direct experimental description of the role of *CREBBP^R1446C^* mutations in lymphoma B-cells, and suggest that *CREBBP* KAT domain mutations may have a potential dominant-negative function on redundant acetyltransferases and thereby drive a more profound molecular phenotype that is associated with a worse patient outcome.

### Synthetic dependence on HDAC3 in *CREBBP*-mutant DLBCL is independent of mutation subtype

Our genomic analysis showed that BCL6 target genes were significantly enriched amongst those with reduced H3K27Ac and gene expression in *CREBBP^R1446C^* cells. We therefore evaluated CREBBP and BCL6 binding over these regions using data from normal germinal center B (GCB)-cells^24^ and found that the regions with *CREBBP^R1446C^* mutation-associated H3K27Ac loss are bound by both proteins (Figure 2A). This suggests that these genes may be counter-regulated by CREBBP and BCL6, the latter of which mediates gene repression via recruitment of HDAC3-containing NCOR/SMRT complexes^3,26^. The epigenetic suppression of gene expression in *CREBBP* mutant cells may therefore be dependent upon HDAC3-mediated suppression of BCL6 target genes. Using a selective HDAC3 inhibitor, BRD3308^27^, we found that *CREBBP*^R1446C^ and *CREBBP^KO^* clones showed greater sensitivity to HDAC3 inhibition compared to isogenic WT controls in cell proliferation assays (Figure 2B). We confirmed this as being an on-target effect of BRD3308 by performing shRNA knock-down of *HDAC3* and observing a greater effect on cell proliferation in *CREBBP*^R1446C^ cells compared to isogenic controls (Figure 2C **and S3**). Moreover, HDAC3 inhibition was able to efficiently promote the accumulation of H3K27Ac in a dose-dependent manner in both *CREBBP^WT^* and *CREBBP^R1446C^* cells, as compared to inactive chemical control compound BRD4097 (Figure 2D). This suggests that the increased sensitivity to HDAC3 inhibition in *CREBBP^R1446C^* cells may be linked with an acquired addiction to an epigenetic change driven by *CREBBP* mutation. We posited that one of these effects may be the suppression of p21 (*CDKN1A*) expression, which is a BCL6 target gene^28^ that has reduced levels of H3K27Ac in both *CREBBP^R1446C^* and *CREBBP^KO^* cells (Figure 2E). In support of this, we observed a marked induction of p21 expression by BRD3308 (Figure 2F) and observed that shRNA-mediated silencing of p21 partially rescued the effect of BRD3308 on cell proliferation (Figure 2G). Therefore *CREBBP* mutations, regardless of type, sensitize cells to the effects of HDAC3 inhibition in part via the induction of p21.

**Figure 2:**
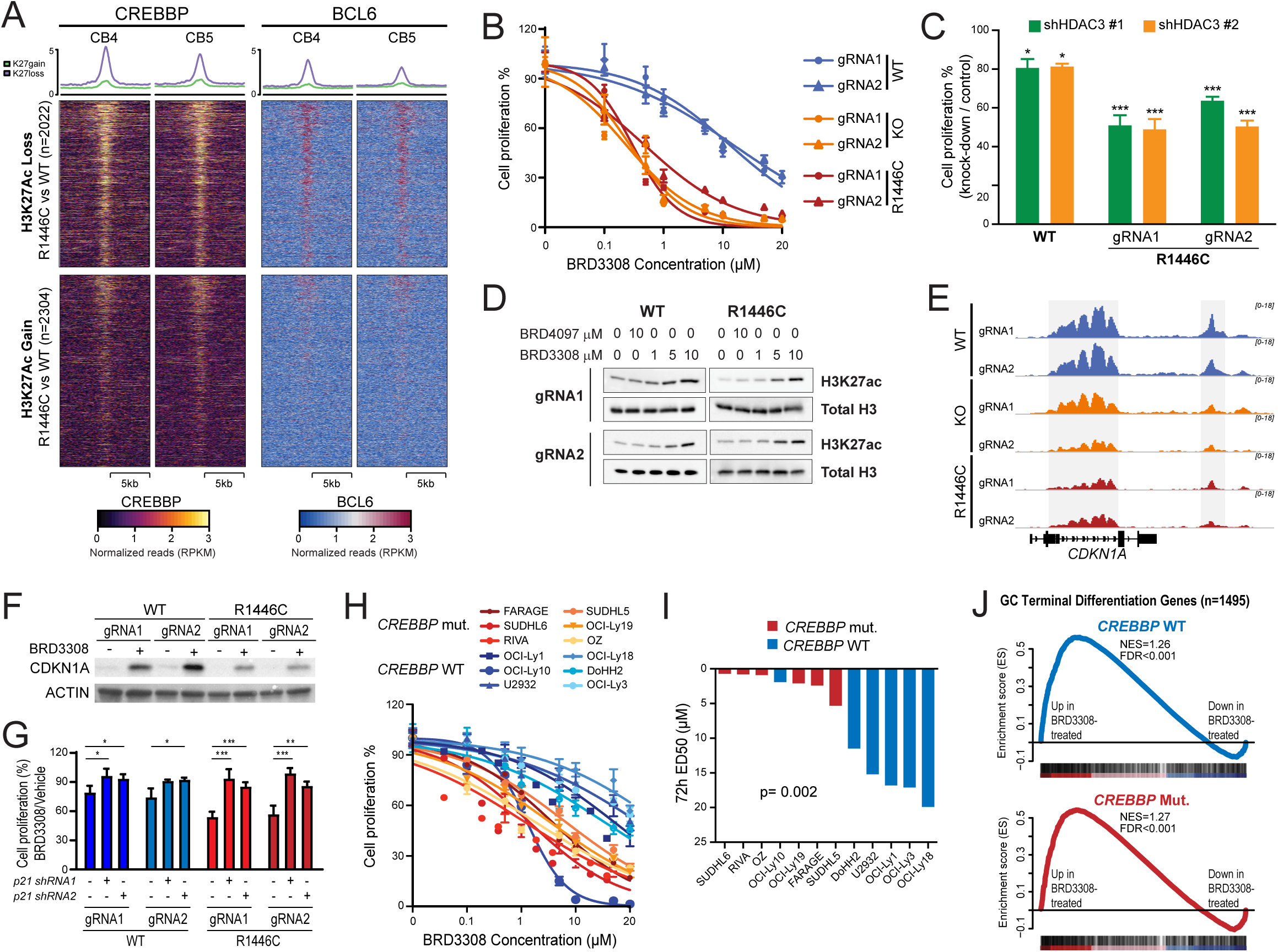
Synthetic dependence upon BCL6 and HDAC3 in CREBBP mutant cells. A) A heat map shows that regions with reduced H3K27Ac in *CREBBP^R1446C^* cells compared to *CREBBP^WT^* cells (above) are bound by both CREBBP and BCL6 in normal germinal center B-cells. This binding is not observed over regions with increased H3K27Ac in mutant cells. **B)** Isogenic *CREBBP^R1446C^* and *CREBBP^KO^* cells have a greater sensitivity to BRD3308, a selective HDAC3 inhibitor, compared to *CREBBP^WT^* cells. **C)** Knock-down of HDAC3 with two unique shRNAs shows a similar preference towards limiting cell proliferation in *CREBBP^R1446C^* cells compared to WT. Data are shown relative to control shRNA in the same cell lines (*P<0.05, ***P<0.001). **D)** Representative western blots show a dose-dependent increase in H3K27Ac in both *CREBBP^WT^* and *CREBBP^R1446C^* cells treated with BRD3308, compared to the control compound BRD4097. **E)** ChIP-seq tracks of H3K27Ac show that *CREBBP^KO^* and *CREBBP^R1446C^* both have reduced levels over the CDKN1A locus compared to isogenic *CREBBP^WT^* cells. Regions that are statistically significant are shaded in grey. **F)** A representative western blot shows that CDKN1A is induced at the protein level by treatment with 10µM BRD3308 in both *CREBBP^WT^* and *CREBBP^R1446C^* cells. **G)** Knock-down of CDKN1A (p21) using two unique shRNAs partially rescued the proliferative arrest of cells treated with BRD3308. This rescue was more significant in *CREBBP* mutant cells compared to wild-type. Data are displayed relative to vehicle-treated cells (*P<0.05, **P<0.01, ***P<0.001). **H)** The difference in sensitivity to BRD3308 between *CREBBP* wild-type (blue) and *CREBBP* mutant (yellow to red) was validated in a large panel of DLBCL cell lines. **I)** The effective dose 50 (ED50) concentrations for each cell line from (H) are shown, colored by *CREBBP* mutation status. The ED50s for *CREBBP* mutant (red) cell lines was significantly lower than that observed for *CREBBP* wild-type cell lines (blue; T-test P=0.002). **J)** Gene set enrichment analysis of Germinal Center Terminal Differentiation signature genes shows that these genes are coordinately induced in both *CREBBP* wild-type (above) and mutant (below) DLBCL cell lines by BRD3308 treatment compared to control.

We aimed to confirm this trend using a larger panel of DLBCL cell lines with either WT (n=6) or mutant *CREBBP* (n=6). This revealed a significantly lower ED50 to BRD3308 in *CREBBP* mutant compared to WT cell lines (p=0.002, Figure 2H-I). We did not observe this trend using the non-specific HDAC inhibitors, Romidepsin and SAHA (**Figure S4**). Notably, none of these cell lines harbor R1446 mutations of *CREBBP*, providing further evidence that sensitivity to HDAC3 selective inhibition is independent of mutation type (i.e. KAT domain missense vs nonsense/frameshift). Furthermore, we also observed the induction of p21 expression and apoptosis in *CREBBP* wild-type cell lines, although to a lesser degree (**Figure S5**). To gain greater insight into this, we performed RNA-sequencing of *CREBBP* wild-type (OCI-Ly1) and mutant (OCI-Ly19, OZ) cell lines treated with BRD3308. Notably, the ability of HDAC3 inhibition to induce the expression of genes involved in the terminal differentiation of B-cells was conserved in both wild-type and mutant cell lines (Figure 2J). Collectively these results are consistent with the role of BCL6 in lymphomas controlling checkpoints, terminal differentiation and other functions^1^ and points towards induction of these transcriptional programs as a potential mechanism of cell death induced by HDAC3-inhibition in both the *CREBBP* WT and mutant settings.

Although targetable by HDAC3 inhibition in wild-type cells, the BCL6-HDAC3 target gene set is more significantly perturbed in the context of *CREBBP* mutation leading to an enhanced cell-intrinsic effect of HDAC3 inhibition.

### HDAC3 inhibition is active against primary human DLBCL

Our observation that HDAC3 inhibition promotes differentiation in both *CREBBP* mutant and wild-type B-cells led us to test its efficacy in primary patient-derived xenograft (PDX) models of DLBCL, for which *CREBBP* mutant tumors were not available. To achieve this, we expanded each tumor *in vivo* and transitioned them to our novel *in vitro* organoid system for exposure to BRD3308^29^. All tumors that were tested showed a dose-dependent reduction in cell viability when cultured with BRD3308, compared to the vehicle control (Figure 3A). We therefore treated 3 of these tumors *in vivo* with either 25mg/kg or 50mg/kg of BRD3308. Quantitative PCR analysis of the treated tumors showed an induction of BCL6 target genes with a role in B-cell terminal differentiation, including *IRF4*, *PRDM1*, *CD138* and *CD40* compared to vehicle-treated tumors (Figure 3B), further supporting the induction of BCL6 target genes as a potential mechanism of cell-autonomous death resulting from HDAC3 inhibition. Furthermore, we also observed a significant reduction in growth of PDX tumors when treated with either dose of BRD3308, thereby providing evidence for *in vivo* efficacy (Figure 3C). Therefore, selective inhibition of HDAC3 may be a rational approach for targeting the aberrant epigenetic silencing of BCL6 target genes in primary human B-cell lymphoma.

**Figure 3:**
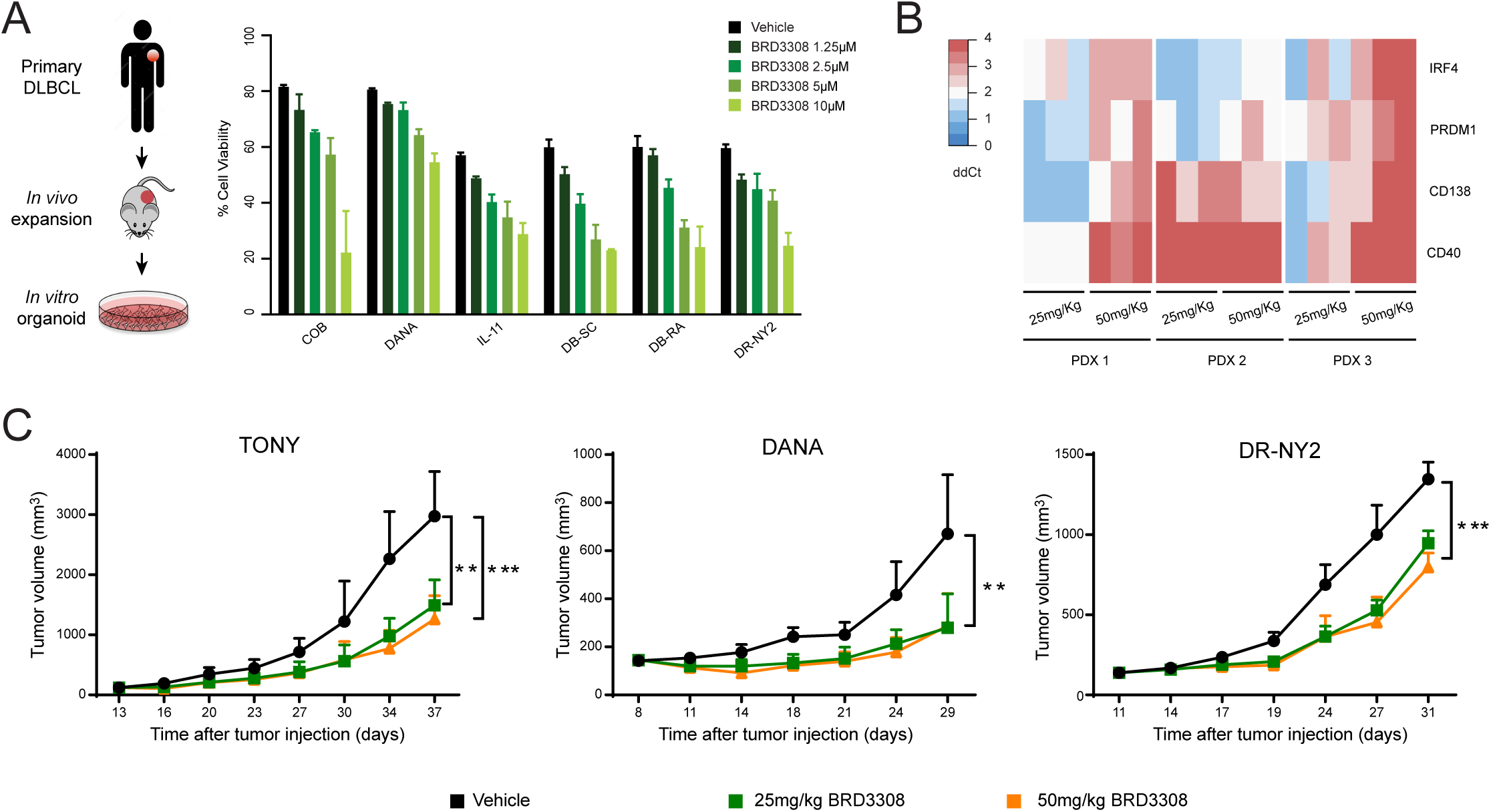
BRD3308 is effective in primary DLBCL. A) The sensitivity of primary DLBCL tumors to BRD3308 was evaluated by expanding them *in vivo*, followed by culture in our *in vitro* organoid model different concentrations of BRD3308. A dose-dependent decrease in cell viability was observed in all 6 tumors with increasing concentrations of BRD3308. B) Treatment of PDX tumors *in vivo* was associated with increased expression of B-cell terminal differentiation genes, as determined by qPCR. The heat map displays the ΔΔC T value of each gene, compared between tumors from treated and untreated mice. **C)** Treatment of 3 unique DLBCL xenograft models *in vivo* with 25mg/kg (green) or 50mg/kg (orange) of BRD3308 significantly reduced tumor growth compared to vehicle (black) (**P<0.01; ***P<0.001).

### Selective inhibition of HDAC3 reverts the molecular phenotype of *CREBBP mutations*

We aimed to take a deeper look at the molecular consequences of HDAC3 inhibition by performing H3K27Ac ChIP-seq and RNA-seq of *CREBBP*^R1446C^ mutant cells after exposure to BRD3308, as compared to negative control compound BRD4097. This showed that selective HDAC3 inhibition promoted the gain of H3K27Ac of a broad number of regions (n=6756, Figure 4A). Strikingly, HDAC3 inhibition either restored or further increased the abundance of H3K27Ac at a large number of sites that became deacetylated in *CREBBP*^R1446C^ and also restored the proper setting for sites where H3K27Ac was increased CRISPR mutated DLBCL cells, consistent with the role of HDAC3 in opposing CREBBP functions (Figure 4B).

**Figure 4:**
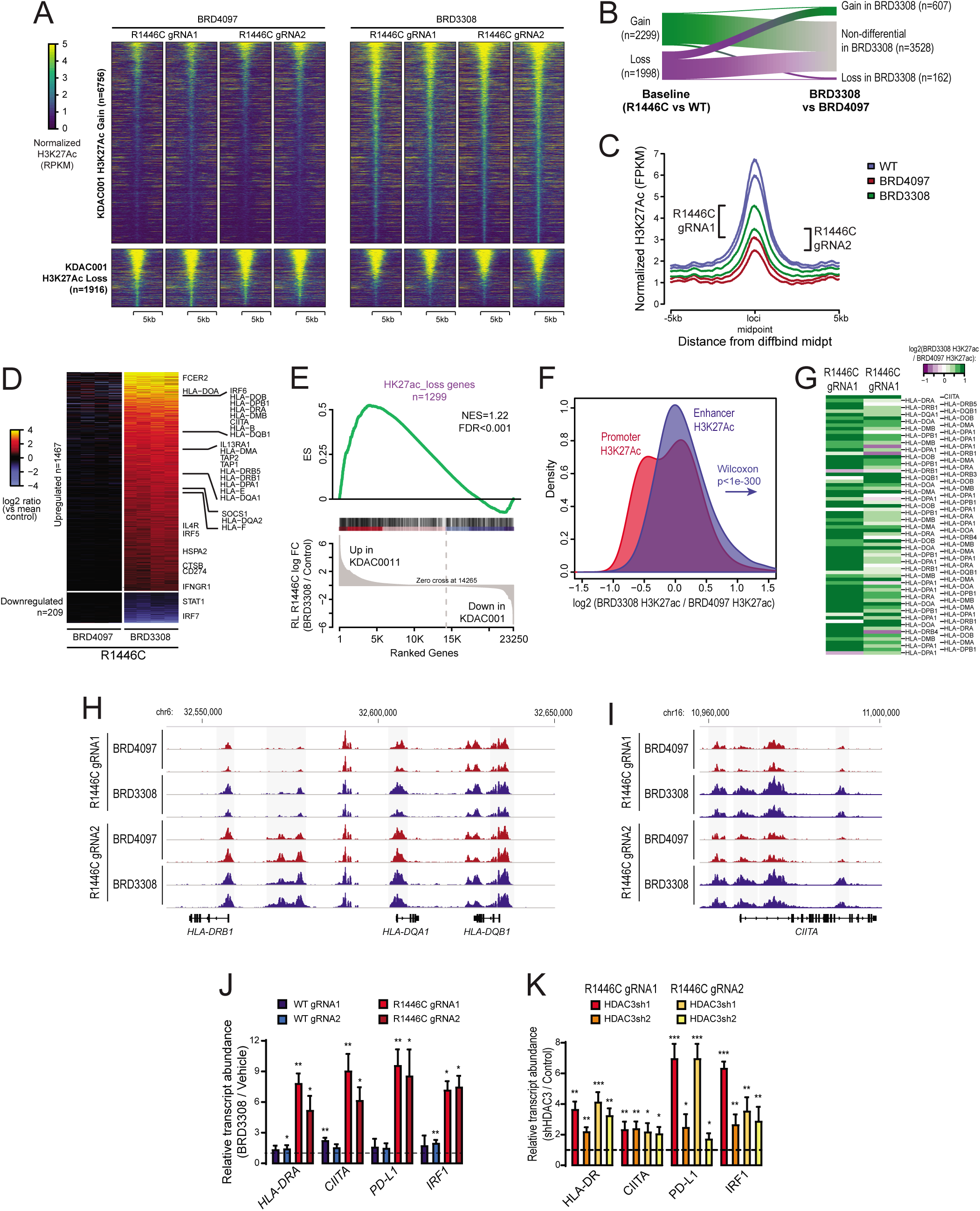
HDAC3 inhibition counteracts the molecular phenotype of *CREBBP* mutation. **A)** A heat map shows the regions with significantly increased (above, n=6756) or decreased (below, n=1916) H3K27Ac in *CREBBP^R1446C^* cells treated with BRD3308 compared to those treated with the control compound, BRD4097. Experimental duplicates are shown for each clone. **B)** A river plot show that a large fraction of the regions with significantly reduced H3K27Ac in *CREBBP^R1446C^* cells compared to *CREBBP^WT^* cells had significantly increased H3K27Ac following BRD3308 treatment. **C)** A density plot shows the regions with reduced H3K27Ac in *CREBBP^R1446C^* compared to *CREBBP^WT^* cells. The level of H3K27Ac over these regions is increased in *CREBBP^R1446C^* cells treated with BRD3308 compared to control (BRD4097), but does not reach the level observed in *CREBBP^WT^* cells. **D)** A heat map shows the genes with increased (above, n=1467) or decreased expression (below, n=209) following BRD3308 treatment. Duplicate experiments are shown for each of the two *CREBBP^R1446C^* clones. Interferon-responsive genes, including those with a role in antigen processing and presentation, can be observed to increase in expression following BRD3308 treatment. **E)** Gene set enrichment analysis shows that the set of genes with reduced H3K27Ac in association with *CREBBP* mutation has coordinately increased expression following BRD3308 treatment. **F)** A density plot illustrates the relative change in promoter (red) and enhancer (blue) H3K27Ac following treatment with BRD3308, with the enhancer distribution being significantly more right-shifted (increased) compared to promoter regions. **G)** A heat map shows the change in H3K27Ac at the promoter regions of MHC class II genes following BRD3308 treatment of *CREBBP^R1446C^* cells, showing a coordinate increase. **H-I)** Regions with significantly increased H3K27Ac (shaded in grey) included those within the MHC class II and *CIITA* gene loci. **J)** The increased expression of candidate genes within the interferon signaling and antigen presentation pathways was confirmed by qPCR. Increased expression was observed in both *CREBBP^WT^* and *CREBBP^R1446C^* cells following BRD3308 treatment, but the level of induction was much higher in *CREBBP^R1446C^* cells. Data are shown relative to vehicle treated cells (T-test *P<0.05, **P<0.01, ***P<0.001). **K)** The on-target role of HDAC3 in the induction of candidate genes was confirmed by shRNA-mediated knock-down of HDAC3 and qPCR analysis of gene expression. Knock-down of HDAC3 was able to induce the expression of all genes, which is shown relative the control shRNA (T-test *P<0.05, **P<0.01, ***P<0.001).

Indeed, a more quantitative analysis indicated HDAC3 inhibition coordinately increased the level of H3K27Ac over the same loci that showed reduced levels in *CREBBP^R1446C^* compared to *CREBBP^WT^* cells (Figure 4C), although this restoration of H3K27Ac was not sufficient to completely revert the epigenomes of *CREBBP^R1446C^* cells to the level that was observed in *CREBBP^WT^* cells (Figure 4C). In line with the role of HDAC3 and BCL6 in transcriptional repression, BRD3308 induced an expression profile that was markedly skewed towards inducing gene upregulation (n=1467 vs 208 genes downregulated; Figure 4D; **Table S5**), including interferon-responsive genes such as antigen presentation machinery and PD-L1 (*CD274*). Notably, the genes with increased expression following HDAC3 inhibition were significantly enriched for those that lose H3K27Ac in *CREBBP^R1446C^* compared to WT (FDR<0.001, Figure 4E), further supporting the conclusion that HDAC3 inhibition directly counteracts changes associated with *CREBBP* mutation. A quantitative assessment of ChIP-seq signal indicated that enhancers manifested greater relative enrichment of H3K27Ac compared to promoters (Wilcoxon P<0.001; Figure 4F), although MHC class II genes also showed coordinate increases in promoter H3K27Ac (Figure 4G). Analysis of critical gene loci in that are deregulated by *CREBBP* mutation, such as MHC class II and CIITA, highlight this induction of H3K27ac (Figure 4H-I). We validated the increased expression of these genes in independent experiments wherein *CREBBP^R1446C^* or isogenic control cells were treated with BRD3308 or vehicle, H3K27Ac was measured by ChIP-qPCR (**Figure S6**), and transcript abundance measured by QPCR (P<0.001, Figure 4J). We further validated that this was an on-target effect of BRD3308 by performing shRNA-mediated knock-down of HDAC3, which resulted in the increased expression of these genes relative to control shRNA (Figure 4K). Together these data indicate that the aberrant mutant-CREBBP epigenetic and transcriptional program can be restored by selective pharmacologic inhibition of HDAC3.

### *CREBBP* mutation dictates the magnitude of induced IFN-responsive and antigen presentation gene expression following HDAC3 inhibition

IFN signaling and antigen presentation genes have not been well investigated as downstream targets of BCL6-HDAC3 complexes, but were enriched in genes that were suppressed by *CREBBP* mutation and restored by HDAC3 inhibition. Given their critical role in anti-tumor immunity, we evaluated whether HDAC3 inhibition may be sufficient to restore or promote the expression of these immune signatures. Using MHC class II protein expression on *CREBBP^R1446C^* mutant cells as a proxy for the CREBBP/BCL6 counter-regulated IFN signaling pathway, we evaluated the activity of HDAC inhibitors for promoting immune response genes. Although HDAC inhibitors with broader specificities were able to induce MHC class II expression, selective inhibition of HDAC3 was sufficient for the robust and maximal (>10-fold) restoration of MHC class II expression in *CREBBP^R1446C^* cells (Figure 5A **and S7**) in line with our observation that these genes are specifically silenced by BCL6/HDAC3 complexes. Furthermore, while some of the less specific HDAC inhibitors were toxic to CD4 and CD8 T-cells, the selective inhibition of HDAC3 was not (**Figure S8**). This suggests that selective HDAC3 inhibition may be capable of eliciting immune responses by on-target on-tumor induction of antigen presentation without on-target off-tumor killing of T-cells.

**Figure 5:**
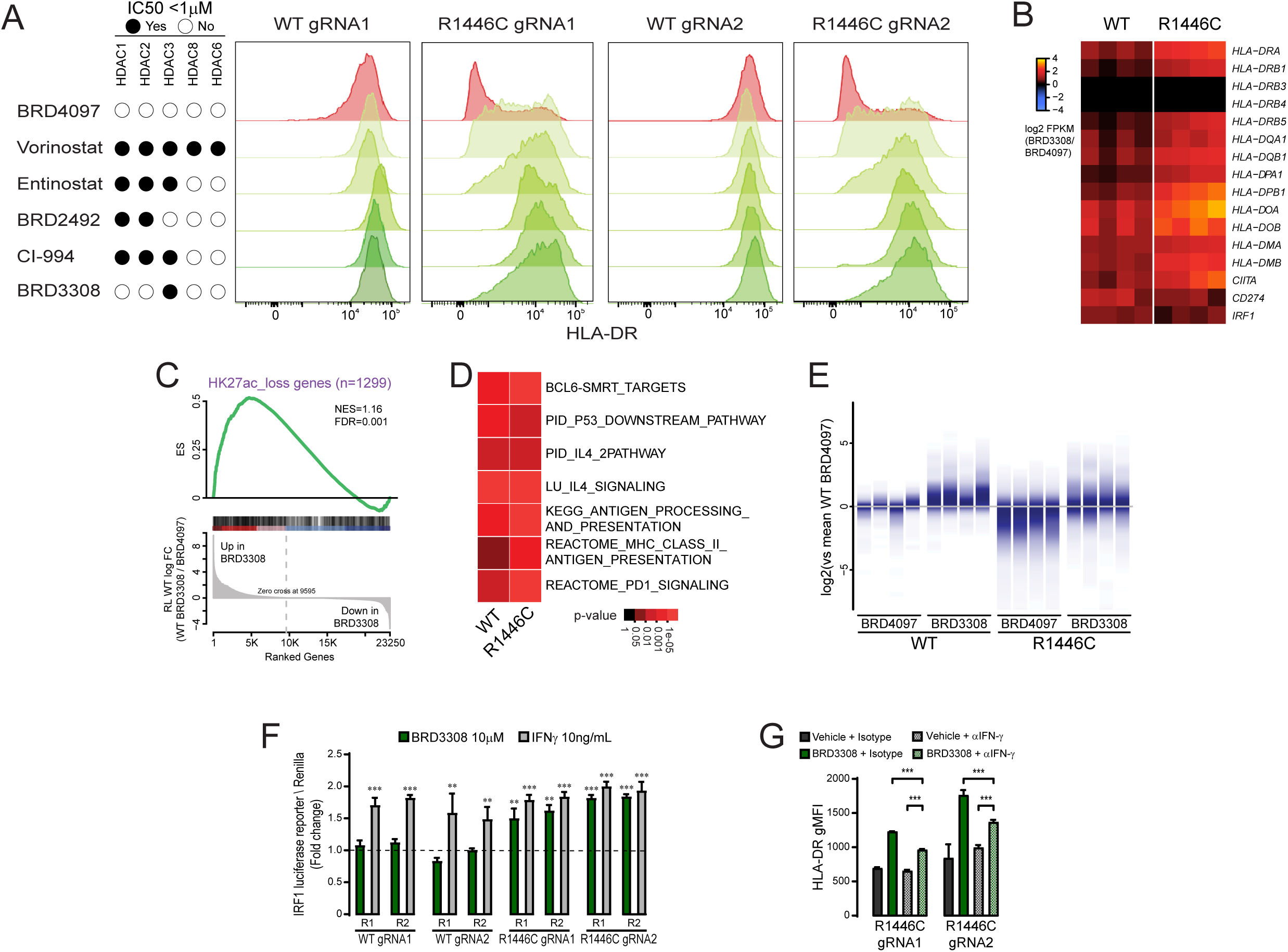
HDAC3 inhibition induces interferon signaling and antigen presentation in both CREBBP wild-type and mutant cells. **A)** Flow cytometry was performed for HLA-DR following exposure to a selection of HDAC inhibitors at 10µM for 72h. This shows that HDAC inhibitors with a range specificities are able to induce MHC class II, but HDAC3 selective inhibition using BRD3308 is sufficient for this effect. **B)** A heat map of interferon responsive and antigen presentation genes from RNA-seq data shows an increased expression in both WT and R1446C cells. Data are normalized to control treated cells from the same experiment. **C)** Gene set enrichment analysis of the genes that have reduced H3K27Ac in *CREBBP^R1446C^* cells shows that the expression of these same genes are coordinately increased by BRD3308 treatment in *CREBBP^WT^* cells. **D)** A heat map of hypergeometric gene set enrichment analysis results of RNA-seq data shows that BRD3308 induces the induction of similar gene set in both *CREBBP^WT^* and *CREBBP^R1446C^* cells. **E)** A density plot, normalized to the mean expression in control (BRD4097)-treated wild-type cells shows the relative expression of the set of genes with reduced H3K27Ac in *CREBBP^R1446C^* cells, normalized to control treated *CREBBP^WT^* cells. This shows that these genes are induced by BRD3308 in *CREBBP^WT^* cells, resulting in expression levels greater than baseline. Further, *CREBBP^R1446C^* cells can be observed to start below baseline, with the induction by BRD3308 resulting in expression levels similar to that observed in control treated *CREBBP^WT^* cells. The 4 samples per condition represent duplicate experiments in each of the two clones for each genotype. **F)** The firefly luciferase luminescence of two unique IRF1 reporters (R1 and R2) is shown, normalized to renilla luciferase from a control vector and shown as fold change compared to untreated cells. *CREBBP^WT^* cells show increased IRF1 activity following IFN-γ treatment (positive control; grey), but not following treatment with BRD3308 (green). In contrast, *CREBBP^R1446C^* cells show increased IRF1 activity following BRD3308 treatment, to a level that is similar to that observed with IFN-γ treatment. (T-test vs control-treated cells, **P<0.01, ***P<0.001). **G)** The role of IFN-γ in inducing MHC class II expression following BRD3308 in *CREBBP^R1446C^* cells was assessed with a blocking experiment. Blocking IFN-γ with a neutralizing antibody (αIFN-γ) significantly reduced the induction of MHC class II, as measured by flow cytometry for HLA-DR, but the induction by BRD3308 with α IFN-γ remained significantly higher than vehicle with α IFN-γ (T-test, ***P<0.001).

Based upon our observations with MHC class II, the magnitude of this induction appeared to be dependent upon the baseline of expression. We therefore hypothesized that *CREBBP* mutation status may determine the magnitude of induction for IFN and antigen presentation pathways, but may not be a prerequisite for this response due to the conserved activity of BCL6/HDAC3 in regulating this axis in WT cells. We comprehensively evaluated this theory using our RNA-seq data from *CREBBP^WT^* and *CREBBP^R1446C^* cells treated with either the HDAC3 inhibitor, BRD3308, or control compound, BRD4097. In parallel to our observations with MHC class II protein expression, BRD3308 treatment coordinately induced the expression of MHC class II and IFN pathway genes in both *CREBBP^WT^* and *CREBBP^R1446C^* cells (Figure 5B). This trend included the majority of genes that were epigenetically suppressed in association with *CREBBP* mutations, resulting in their significant increase in expression in *CREBBP^WT^* cells (Figure 5C) similar to our observations from *CREBBP^R1446C^* cells (Figure 4F). Moreover, we further observed similar enrichment of transcriptionally-induced pathways in both *CREBBP^WT^* and *CREBBP^R1446C^* cells that included the same pathways that were suppressed by *CREBBP* mutation (Figure 5D; **Table S6 and S9**). Consistent with the almost exclusive function of HDAC3 as a BCL6 corepressor in GC B-cells^3^, and the importance of BCL6 activity in both *CREBBP* WT and mutant cells, we observed highly significant enrichment for genes regulated by BCL6-SMRT complexes among genes with induced H3K27Ac and expression after BRD3308 treatment (Figure 5D). Also significantly enriched were canonical BCL6 target gene sets such as p53 regulated genes, and signaling through B-cell receptor, CD40 and cytokines like IL4 and IL10. Finally we observed significant enrichment for BCL6 target gene sets linked to IFN signaling, antigen presentation via MHC class II and PD1 signaling.

Although there were conserved patterns of gene activation in both *CREBBP^WT^* and *CREBBP^R1446C^* cells, we observed that the magnitude of this induction was greatest in *CREBBP^R1446C^* cells which started from a lower baseline of expression linked to mutation-associated epigenetic suppression (Figure 5E). We identified IRF1 as a BCL6-regulated transcription factor that is critical for interferon responses, and is induced by HDAC3 inhibition *CREBBP^WT^* and *CREBBP^R1446C^* cells (Figure 5B). We therefore hypothesized that IRF1 may contribute to the different magnitude of induction in MHC class II genes following HDAC3 inhibition in these two genetic contexts. We investigated this by measuring IRF1 activity in a luciferase reporter assay in *CREBBP^WT^* and *CREBBP^R1446C^* mutant cells. Treatment of *CREBBP*^R1446C^ cells with BRD3308 led to a significant increase of IRF1 activity that was similar in magnitude to that induced by IFN-γ treatment (Figure 5F). In contrast, this effect was not observed in isogenic *CREBBP^WT^* control cells following HDAC3 inhibition, despite these cells showing similarly increased IRF1 activity in response to IFN-γ treatment. Exposing *CREBBP*^R1446C^ cells to IFN-γ neutralizing antibodies only partially ameliorated MHC class II induction after BRD3308 in *CREBBP* mutant cells, but did not completely eliminate the effect (Figure 5G). This suggests that preferential induction of antigen presentation by HDAC3 inhibition in *CREBBP* mutant cells likely depends on a combination of mechanisms, including direct BCL6 repression of these genes as well as secondary induction through IFN-γ signaling (which is also directly regulated by BCL6).

### HDAC3 inhibition restores MHC class II expression in human DLBCL cell lines and patient specimens

The frequency of MHC class II loss in DLBCL exceeds the frequency of *CREBBP* mutations in this disease^12,21^, through unknown mechanisms. The ability of HDAC3 inhibition to induce MHC class II expression in *CREBBP* WT DLBCL cells may therefore have potential broad implications for immunotherapy. Using RNA-sequencing data from WT (OCI-Ly1) and mutant (OCI-Ly19 and OZ) DLBCL cell lines, we confirmed that the set of genes induced by HDAC3 inhibition was largely conserved in both contexts (Figure 6A). In a broader panel of *CREBBP* WT and mutant cell lines, we observed that a core set of genes including *HLA-DR*, *CIITA* and *PD-L1* had consistently higher expression in BRD3308-treated cells compared to the matched control, but with a higher magnitude of induction in *CREBBP* mutant cell lines (Figure 6B). This trend was also observed by flow cytometry for MHC class II, which extends upon our observations in CRISPR/Cas9-modified cells by showing a reproducible increase in expression in a larger panel of DLBCL cell lines, and a higher magnitude of induction in *CREBBP* mutant cell lines (Figure 6C). These observations validate those made in CRISPR/Cas9-modified cells, showing that antigen presentation can be promoted by HDAC3 inhibition in *CREBBP* WT cell-lines. Although *CREBBP* mutant PDX tumors were not available for analysis, we observed that one of our PDX tumors was MHC class II negative at baseline despite being *CREBBP* WT (Figure 6D). The *in vivo* treatment of this PDX with the HDAC3 inhibitor, BRD3308, was sufficient to restore MHC class II expression in this tumor in a dose-dependent manner (Figure 6D). This effect was also observed when untreated cells from this PDX were cultured in our organoid system in the presence of BRD3308 vs. vehicle, and the expression of HLA-DR and PD-L1 assessed by qPCR (Figure 6E).

**Figure 6:**
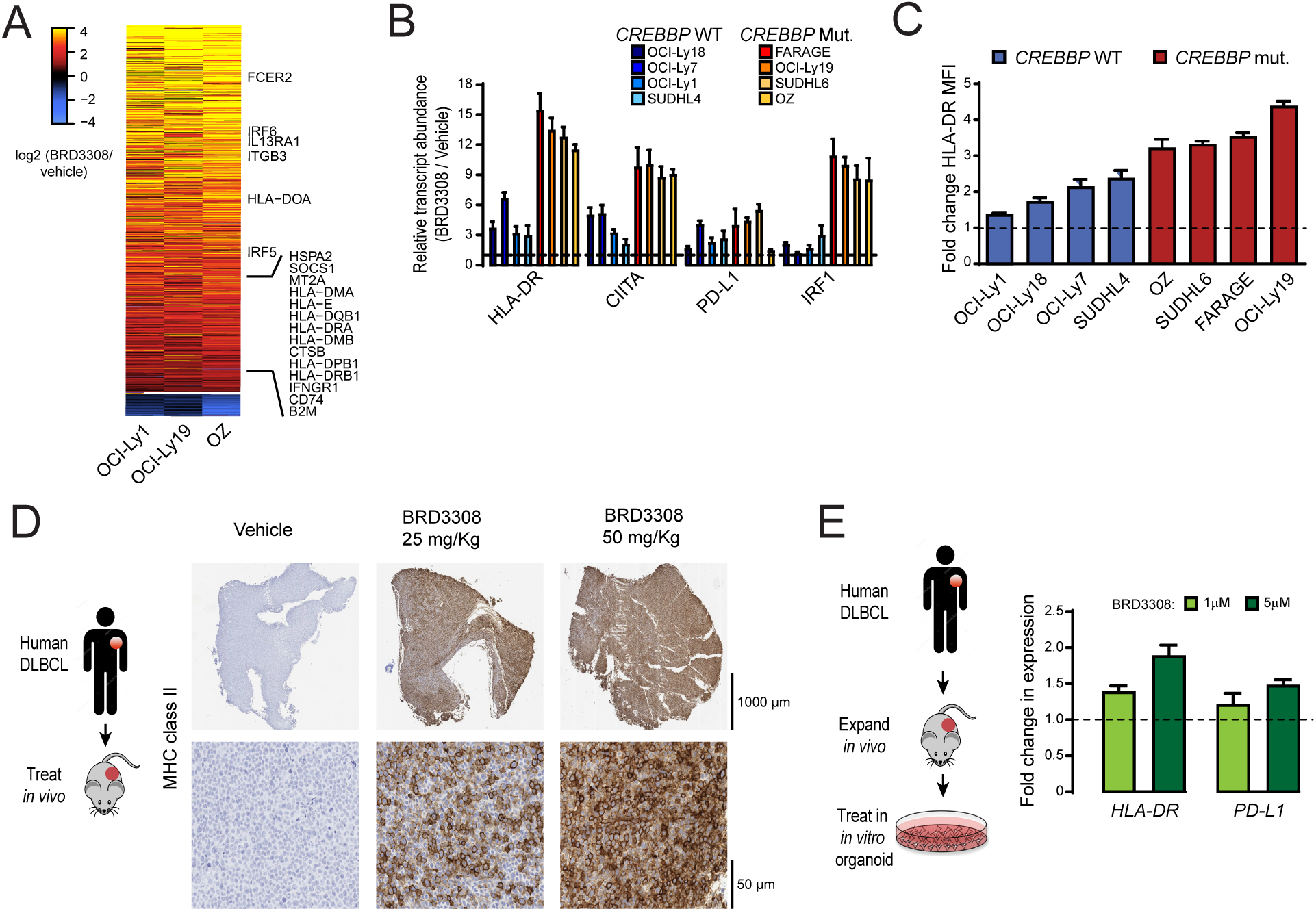
Induction of interferon-responsive and antigen presentation genes in DLBCL cell lines and patient-derived xenograft. **A)** A heat map shows significantly up-regulated (above) and down-regulated (below) genes in BRD3308-treated DLBCL cell lines that are *CREBBP* wild-type (OCI-Ly1) or mutant (OCI-Ly19 and OZ), expressed as a log2 ratio to vehicle control treated cells. The observed changes were consistent between wild-type and mutant cell lines, and included up-regulation of interferon-responsive and antigen presentation genes. **B)** qPCR was used to validate the gene expression changes of select interferon-responsive genes following BRD3308 treatment across an extended panel of *CREBBP* wild-type and mutant DLBCL cell lines. These genes were uniformly increased in both genetic contexts, but with a higher magnitude of increase in *CREBBP* mutant cell lines. **C)** The induction of MHC class II expression by BRD3308 was measured in an extended panel of DLBCL cell lines by flow cytometry. Data are plotted as a fold-change of the mean fluorescence intensity (MFI) of HLA-DR in BRD3308-treated vs control-treated cells. We observed uniformly increased MHC class II expression in all cell lines, but with higher magnitude in *CREBBP* mutants. **D)** An MHC class II-negative DLBCL patient derived xenograft model was treated *in vivo* with either 25mg/kg or 50mg/kg of BRD3308. Immunohistochemical staining was performed for MHC class II, revealing a robust induction of MHC class II expression that was relative to the dose of treatment. **E)** The MHC class II negative PDX was expanded *in vivo* and then treated with BRD3308 in an *in vitro* organoid culture. The expression of the interferon-responsive genes HLA-DR and PD-L1 (*CD274*) were measured by qPCR and found to increase in a dose-dependent manner. Bars represent the fold-change compared to vehicle control +/- S.D..

Together, these results show that selective inhibition of HDAC3 using BRD3308 can promote the expression of IFN and antigen presentation pathway genes in both the *CREBBP* WT and mutant settings. However, the magnitude of induction is greatest in *CREBBP* mutant cells owing, in part, to the preferential induction of IRF1 activity in these cells.

### HDAC3 inhibition drives T-cell activation and antigen-dependent immune responses

Interferon signaling and antigen presentation are central to anti-tumor immune responses. We therefore investigated whether HDAC3 inhibition could promote antigen-dependent anti-tumor immunity. For this experiment we implanted OCI-Ly18 DLBCL cells into NSG mice and once tumors formed we injected human peripheral blood mononuclear cells (PBMCs) including T-cells to educate them to tumor antigens. After *in vivo* priming, tumor infiltrating lymphocytes (TILs) were then co-cultured *in vitro* with OCI-Ly18 cells that were pre-treated for 72h with increasing concentrations of BRD3308 to assess the effects on T-cell activation and tumor cell killing (Figure 7A). The DLBCL cells that were epigenetically primed for antigen presentation by BRD3308 increased the activation of CD4 T-cells, as determined by CD69 expression (Figure 7B). These tumor-educated T-cells also produced higher levels of IFN-γ when co-cultured with OCI-Ly18 cells that had been primed with HDAC3 inhibitor (Figure 7C **and S9**). As in prior experiments, we observed cell-intrinsic effects of BRD3308 on OCI-Ly18 cells in the absence of TILs, resulting in declining cell viability with higher concentrations of the inhibitor. However, the effects of BRD3308 were markedly increased in the presence of TILs, consistent with T-cell-directed killing of the tumor cells (Figure 7D). To confirm that this killing was dependent on MHC:TCR interactions we also performed this experiment in the presence of blocking antibodies for MHC class I, MHC class II or both. Blocking one or the other of MHC class I or II rescued some of the cytotoxicity observed in this assay, but blocking both MHC class I and class II completely abrogated the TIL-associated effect (Figure 7D). These data therefore show that HDAC3 inhibition can potentiate anti-tumor immune responses that are likely to be antigen-dependent because they are driven by MHC:TCR interactions.

**Figure 7:**
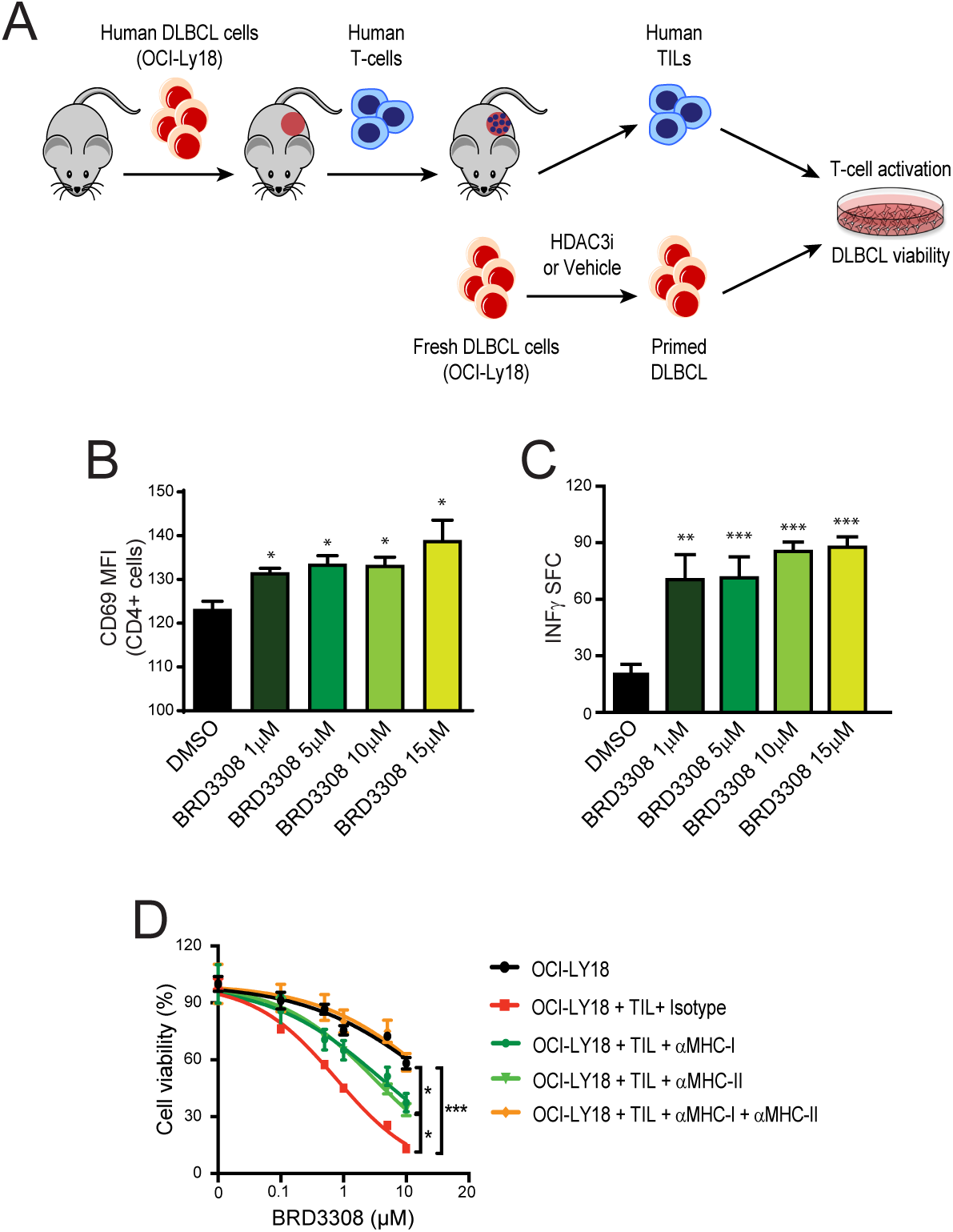
HDAC3 inhibition induces antigen-dependent immune responses. **A)** A schematic of the generation of antigen-specific T-cells and epigenetic priming of DLBCL cells. A human DLBCL cell line (OCI-Ly18) was engrafted into immunodeficient mice and allowed to establish. Human T-cells were then engrafted, allowing them to become educated to the tumor antigens prior to harvesting of the tumor-infiltrating T-cell (TIL) fraction. These TILs were cultured with fresh DLBCL cells that had been epigenetically primed with different concentrations of BRD3308, and the cell viability of the DLBCL cells measured after 72h. **B)** TIL and DLBCL co-culture resulted in activation of the CD4 T-cells in a dose-dependent manner, as measured by flow cytometry for the CD69 activation marker. Data represent the fold change in CD69 expression compared to vehicle treated DLBCL cells. **C)** The production of IFN-γ was measured by ELISPOT and found to increase in cultures with epigenetically-primed DLBCL cells. (T-test vs DMSO control, **P<0.01, ***P<0.001) **D)** The cell viability of DLBCL cells in TIL co-culture experiments was measured by CellTiterBlue assay. Treatment with BRD3308 resulted in some cell killing through cell-intrinsic mechanisms in the absence of TILs (black). The addition of TILs at a 1:1 ratio led to a significant increase in cell death of the DLBCL cells. This was partially reduced by blocking of either MHC class I or MHC class II using neutralizing antibodies. Blocking of MHC class I and class II together completely eliminated the TIL-associated increase in cell death, suggesting that killing was mediated through MHC:TCR interactions.

## DISCUSSION

Precision medicine and immunotherapy have led to significant breakthroughs in a variety of cancers, but have lacked success in B-cell lymphoma. For precision medicine, this is largely due to a paucity of ‘actionable’ genetic alterations or, rather, the lack of current therapeutic avenues to target the mutations that have been defined as being important for disease biology. For immunotherapy, the mechanisms driving lack of response or early progression are not well understood, but are likely to be underpinned by the complex immune microenvironment and genetic alterations that drive immune escape. The exception to both of these statements is the use of PD-1 neutralizing antibodies in classical Hodgkin lymphoma, which opposes genetically-driven immune suppression by the malignant Reed-Sternberg cells through DNA copy number gain of the PD-L1 locus^30^ and elicits responses in the majority of patients^31^. This stands as an example of the potential success that could be achieved by the characterization and rational therapeutic targeting of genetic alterations and/or the neutralization of immune escape mechanisms. However, we are not yet able to successfully target some of the most frequently mutated genes or overcome the barriers of inadequate response to immunotherapy in the most common subtypes of B-cell lymphoma. These are important areas of need if we hope to further improve the outcomes of these patients.

The *CREBBP* gene is mutated in ~15% of DLBCL^21^ and ~66% of FL^6^, and is therefore a potentially high-yield target for precision therapeutic approaches. Our use of CRISPR/Cas9 gene-editing to generate isogenic lymphoma cell lines that differ only in their *CREBBP* mutation status allowed us to perform detailed characterization of the epigenetic and transcriptional consequences of these mutations. Using this approach we identified for the first time functional differences between the most frequent KAT domain point mutation, R1446C, and frameshift mutations that result in KO. Although similar regions of the genome showed reduced H3K27Ac in R1446C and KO mutants compared to isogenic WT controls, the magnitude of these changes were markedly reduced in *CREBBP* KO. This suggests that R1446C mutations of *CREBBP* may act in a dominant-negative fashion by preventing the participation of redundant acetyltransferases in transcriptional activating complexes. But in the setting of *CREBBP* nonsense/frameshift mutations, acetyltransferases such as EP300 may compensate for the loss of CREBBP protein. This is an important observation because, to date, *CREBBP* mutations have only been modeled in mice using conditional knock-out or knock-down of the gene, which we may now expect to results in a less severe phenotype compared to KAT domain point mutations.

Despite differences in the magnitude of molecular changes between R1446C and KO cells, these mutations yielded a similar degree of synthetic vulnerability to HDAC3 inhibition. This was likely driven by the increased suppression of BCL6 target genes that we observed in both contexts, including *CDKN1A* (p21). This gene has also been highlighted as a critical nexus in the oncogenic potential of *EZH2*^32^, which cooperates with BCL6 to silence gene expression^4^. One of the important mechanisms for BCL6-mediated gene silencing is through the recruitment of HDAC3 as part of the NCOR/SMRT complex^3^, thereby highlighting HDAC3 inhibition as a rational therapeutic avenue for counteracting BCL6 activity. Virtually all of the HDAC3 corepressor complexes present in DLBCL cells are bound with BCL6, suggesting that HDAC3 inhibitors effect is largely explained by their depression of the subset of BCL6 target genes regulated through this mechanism^3^. Importantly, as a variety of GCB-derived malignancies rely on BCL6 function independently of *CREBBP* or *EZH2* mutation^33,34^, opposing this function through HDAC3 inhibition may also be effective in tumors without these genetic alterations. We have shown preliminary evidence in the primary setting using DLBCL PDX models, for which *CREBBP* mutant tumors were not currently available. However, we would expect to observe greater efficacy in the *CREBBP* mutant setting, as we have shown in CRISPR/Cas9-edited cells and DLBCL cell lines.

One of the important transcriptional programs that is regulated by BCL6 is the interferon signaling pathway^3^, which we observed to be significantly repressed in *CREBBP* mutant cells. It has been also long been recognized that IFN-γ cooperates with lymphocytes to prevent cancer development^35^. Interferon signaling supports T-cell driven anti-tumor immunity via a variety of mechanisms, including the induction of antigen presentation on MHC class II^36^. We have shown that the selective inhibition of HDAC3 is sufficient for broadly restoring the reduced H3K27Ac and gene expression that is associated with *CREBBP* mutation, including the interferon signaling and antigen presentation programs. This was in part driven by the increased production of IFN-γ following HDAC3 inhibition, but also via the induced expression and activity of the IRF1 transcription factor. Together, these factors lead to a robust restoration of MHC class II expression on *CREBBP* mutant cells and drove dose-dependent potentiation of anti-tumor T-cell responses. However, we also noted that the same molecular signature is promoted by HDAC3 inhibition in *CREBBP* wild-type cells, also with an associated increase of MHC class II expression. As with the cell-intrinsic effects of HDAC3 inhibition, the effects on immune interactions may therefore be active in both *CREBBP* wild-type and *CREBBP* mutant cells, as a result of the conserved molecular circuitry controlling these pathways in each genetic context.

A variety of HDAC inhibitors were capable of restoring antigen presentation in our models and clinical responses are observed with these agents in relapsed/refractory FL and DLBCL patients. However, grade 3-4 hematological toxicities such as thrombocytopenia, anemia and neutropenia were frequent, and the responses tended not to be durable^37,38^. We posit that specific inhibition of HDAC3 may be accompanied by reduced toxicity, as HDAC1 and HDAC2 have important roles in hematopoiesis^39^ and avoiding the inhibition of these HDACs may therefore avoid the undesired hematological effects associated with pan-HDAC inhibition. In addition, we speculate that the clinically-observed lack of durability may be the result of adaptive immune suppression through mechanisms such as PD-L1 expression, which dampens T-cell responses through the PD-1 receptor^40^, as well as direct toxicity of pan-HDAC inhibitors to T-cells. We found evidence for adaptive immune suppression within our model systems, showing that HDAC3 inhibition leads to increased IFN-γ production and the upregulation of *PD-L1* expression. This is in line with recent observations that PD-L1 (*CD274*) is a BCL6-supressed gene^41^, and a well-characterized role for PD-L1 as an IFN-γ-responsive gene^40^. In other cancers in which a florid antigen-driven immune response and concomitant adaptive immune suppression via PD-L1 exist within the tumor microenvironment, blockade of the PD-1 receptor is an effective therapeutic strategy^13,15^. In support of this, recent studies have shown that the efficacy of PD-1 blockade is inextricably linked with the existence of an interferon-driven immune response and expression of MHC class II^13,15^. Together, these observations suggest that the greatest potentiation of anti-tumor immunity in GCB-derived malignancies may be achieved through stimulation of interferon signaling and MHC class II expression by HDAC3 inhibition, in combination with the inhibition of adaptive immune suppression using PD-L1/PD-1 neutralizing antibodies. However, this concept requires further exploration in future studies.

In conclusion, this work defines a molecular circuit that controls the survival and differentiation of lymphoma B-cells and their interaction with T-cells. This circuit is counter-regulated by CREBBP and BCL6 and can be pushed towards promoting tumor cell death and anti-tumor immunity via selective inhibition of HDAC3. This highlights HDAC3 inhibition as an attractive therapeutic avenue, which may be broadly active in FL and DLBCL due to the near-ubiquitous role of BCL6, but which may have enhanced potency in *CREBBP* mutant tumors.

## ACKNOWLEDGEMENTS

This work was supported by R01 CA201380 (MRG), R01 CA055349 (DAS), U54 OD020355 01 (ES and GI), NCI SPORE P50 CA192937 (AY), the MD Anderso Cancer Center (P30 CA016672) and Memorial Sloan Kettering Cancer Center (P30 CA008748) NCI CORE Grants, the Chemotherapy Foundation (AM), the Star Cancer Consortium (AM), and the Jaime Erin Follicular Lymphoma Research Consortium (AM, MRG, SN).

